# Ventral pallidal perineuronal nets regulate opioid relapse

**DOI:** 10.64898/2026.01.21.700926

**Authors:** Margareth Nogueira, Giuseppe Giannotti, Carley N. Miller, Savanna D. Guaderrama, Nathaniel P. Kregar, Brandi Wiedmeyer, Nicholas Fayette, Jasper A. Heinsbroek

## Abstract

Opioid use disorder remains a major health challenge worldwide. Neuronal activity in the ventral pallidum (VP) regulates opioid reward and relapse to opioid seeking but the underlying cellular mechanisms remain largely unknown. A sizable population of VP neurons previously linked to drug relapse expresses the calcium binding protein parvalbumin (VP_PV_). Across the brain parvalbumin neurons are often ensheathed by perineuronal nets (PNNs), specialized extracellular structures that regulate intrinsic activity and constrain synaptic plasticity onto these neurons. The VP contains high levels of PNNs but the role of these structures in the neurophysiology of VP_PV_ neurons and in relapse to opioid seeking has not been studied. To investigate whether VP PNNs are altered by opioid exposure, male and female mice were trained to self-administer intravenous heroin. We found that heroin increased the density of PNNs in the VP, and that an intracranial microinfusion of the PNN-degrading enzyme, chondroitinase ABC, prevented cue-induced reinstatement of heroin seeking. VP PNN depletion also reduced the intrinsic excitability of VP_PV_ neurons, potentiated inhibitory synaptic inputs onto these cells, and diminished Fos expression in VP_PV_ neurons following reinstatement. The suppressive effect of VP PNN depletion on heroin seeking was rescued by chemogenetic activation of VP_PV_ neurons and mimicked by chemogenetic VP_PV_ neuron inhibition. Taken together, our results identify VP_PV_ neurons and their associated PNNs as critical drivers of opioid seeking. Given the key role of PNNs in regulating neural plasticity and memory processes, targeting PNNs in the VP could provide a useful novel therapeutic avenue for treating persistent craving and relapse in opioid use disorder.

## Introduction

The use of opioids and opioid-related overdose deaths have increased drastically over the last decade. Persistent opioid use can lead to opioid use disorder (OUD), which is characterized by dependence, continued use despite negative consequences, and often, by recurrent cycles of withdrawal and relapse (Koob & Volkow, 2016). Opioids produce long-lasting neurobiological changes in the interconnected nuclei of the ventral basal ganglia that are associated with persistent drug craving and relapse risk (Heinsbroek et al., 2021; Kruyer et al., 2020). The ventral pallidum (VP) is a key node within this network critical for integrating reward-related information to direct motivated behavior (Kalivas, 2009; Root et al., 2015; Soares-Cunha & Heinsbroek, 2023). Importantly, neuronal activity in the VP is critical for relapse to opioid seeking (Farrell et al., 2022; Rogers et al., 2008).

Exposure to drugs of abuse persistently alters the functioning of VP neurons, but the mechanisms whereby chronic drug use produces lasting changes in these neurons remain largely unknown (Root et al., 2015; Soares-Cunha & Heinsbroek, 2023). A substantial population of VP neurons linked to drug relapse is characterized by the expression of the calcium-binding protein parvalbumin (VP_PV_) (Celio, 1990; Gritti et al., 2003; Prasad et al., 2020). Importantly, persistent functional changes have been reported in VP_PV_ neurons in response to aversive experiences that alter motivation and hedonic states (Knowland et al., 2017), linking changes in their function to long-lasting behavioral adaptations. Although VP_PV_ neurons have been implicated in relapse to drug seeking (Prasad et al., 2020), it remains unclear how exposure to drugs of abuse could elicit persistent maladaptive plasticity in this population.

Across the brain many PV neurons are ensheathed by perineuronal nets (PNN), specialized extracellular matrix structures that regulate neuronal activity and that constrain neuroplasticity by limiting synaptic and astroglial connections onto these cells (Lupori et al., 2023; Reichelt et al., 2019; Tewari et al., 2024; Wingert & Sorg, 2021). PNNs also support the fast-spiking characteristics of PV neurons by trapping cations and signaling molecules near the soma and by stabilizing synaptic inputs (Hartig et al., 1999; Morawski et al., 2015; Song & Dityatev, 2018). In many systems the maturation of PNNs mediates the closure of critical periods of neurodevelopment, and their ability to functionally stabilize the neurons they surround has been suggested to play a key role in long-term memory storage (Fawcett et al., 2022; Tsien, 2013).

PNNs are implicated in the maladaptive plasticity that underlies persistent drug memories in substance use disorders. Drugs of abuse alter the density of PNNs in many brain regions, and their removal by enzymatic digestion perturbs drug memories (Blacktop & Sorg, 2019; Chen et al., 2015; Chen & Lasek, 2020; Slaker et al., 2015; Van den Oever et al., 2010). PNN depletion can also induce plasticity and facilitate new learning, such as the extinction of drug seeking behavior (Xue et al., 2014). Collectively, these findings strongly implicate PNNs in the formation and maintenance of persistent drug memories (Fawcett et al., 2022).

Prior work has shown a dense expression of PNNs in the VP, but their function in this structure has not yet been explored (Brauer et al., 1993; Seeger et al., 1994). We hypothesize that VP PNNs contribute to opioid seeking by stabilizing VP_PV_ neurons and by increasing their intrinsic activity to drive motivation for opioids. To test this, we investigated whether changes in the density of VP PNNs occur following intravenous heroin or saline self-administration. We then employed the PNN ablating enzyme chondroitinase ABC (ChABC) to investigate the role of VP PNNs in heroin seeking and in the neurophysiology of VP_PV_ neurons. Finally, we used chemogenetics to test whether stimulating VP_PV_ neurons would be sufficient to restore drug seeking following PNN depletion, and whether inhibiting these neurons would recapitulate the effects of PNN depletion and suppress heroin seeking. Results from this multifaceted approach firmly implicate VP PNNs and the VP_PV_ neurons they surround in relapse to opioid seeking.

## Results

### Perineuronal nets are widely expressed in the VP and preferentially localize to VP_PV_ neurons

Prior studies have reported high levels of PNNs in the VP with mixed results on whether PNNs in the VP surround VP_PV_ neurons (Adams et al., 2001; Brauer et al., 1993; Lupori et al., 2023; Seeger et al., 1994). We reasoned that part of this discrepancy could be mediated by poor antigen and lectin availability due to high levels of neuropil in the VP and observed that the labeling of PNNs with the lectin binding label wisteria floribunda agglutin (WFA) was substantially improved by adding sodium nitrate and heparin to our exsanguination solution prior to the transcardial formalin fixation of brain tissue (Marchant et al., 2009). This approach also allowed us to clearly visualize PNNs in the VP and nucleus accumbens (Figure 1a) where mixed findings have been reported regarding their presence (Bertolotto et al., 1996; Hazlett et al., 2024; Lee et al., 2012; Seeger et al., 1994). Next, we quantified the co-localization between PNNs and PV immunohistochemistry in the VP using confocal microscopy. This analysis showed that most neurons surrounded by a PNN are PV positive (Figure 1b; paired t-test comparing between PV positive and PV negative neurons surrounded by a PNN: t_(5)_ = 23.87, p = 2.404 × 10^-6^) and that over two-thirds of VP_PV_ neurons are surrounded by a PNN (paired t-test comparing between PV neurons with or without a PNN: t_(5)_ = 5.370, p = 2.404 × 10^-4^).

**Figure 1.**
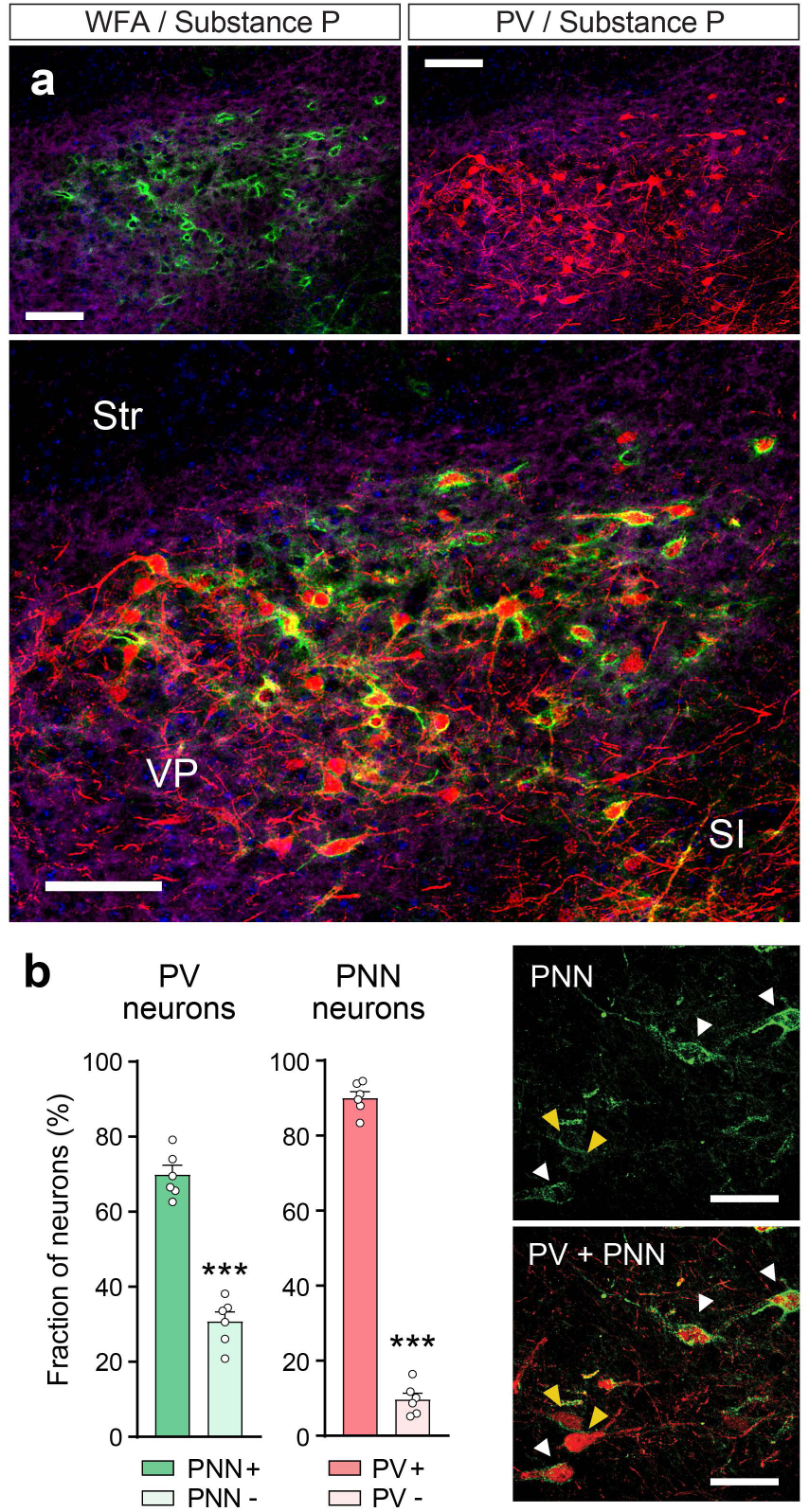
PNNs in the VP preferentially surround VP_PV_ neurons. (**a**) Micrographs showing PNN labeling (top left; green) and PV labeling (top right; red) in the VP, and the preferential localization of PNNs to VP_PV_ neurons. Tissue was counterstained with substance P to delineate the boundaries of the VP (purple). Str = striatum. SI = substantia innominata. Scale bar = 100 μm. (**b**) Over two thirds of VP_PV_ neurons are surrounded by a PNN (green), and the majority of PNNs in the VP ensheathe PV neurons (red). Inserts on the right show VP_PV_ neurons surrounded by dense (white arrow) and faint PNNs (yellow arrow). Scale bar = 50 μm. Symbols in bars represent individual mice (n = 6). *** p < 0.001. Data are shown as mean ± s.e.m.

### Heroin self-administration increases PNN density in the VP

To examine the persistent effects of a prior history of heroin self-administration on PNNs in the VP, we surgically implanted mice with intravenous jugular vein catheters and trained them to self-administer heroin or saline. Mice responded with significantly more active but not inactive nose pokes for heroin compared to saline (Figure 2a; RM-ANOVA main effect of drug F_(1,10)_ = 198.8, p = 6.329 × 10^-8^). After 14 days of intravenous heroin or saline self-administration and 7 days of extinction (Figure 2b), mice were transcardially perfused for PV immunohistochemistry and WFA-lectin labeling (Figure 2c). Heroin self-administration produced a significant increase in the density of perineuronal nets across the VP compared to saline controls (Figure 2d; Mann-Whitney U = 5, p = 0.048), but did not significantly alter PV expression. These data indicate that a prior history of heroin self-administration increases the density of PNNs and suggests that this structural change may alter the functioning of VP_PV_ neurons to promote heroin seeking.

**Figure 2.**
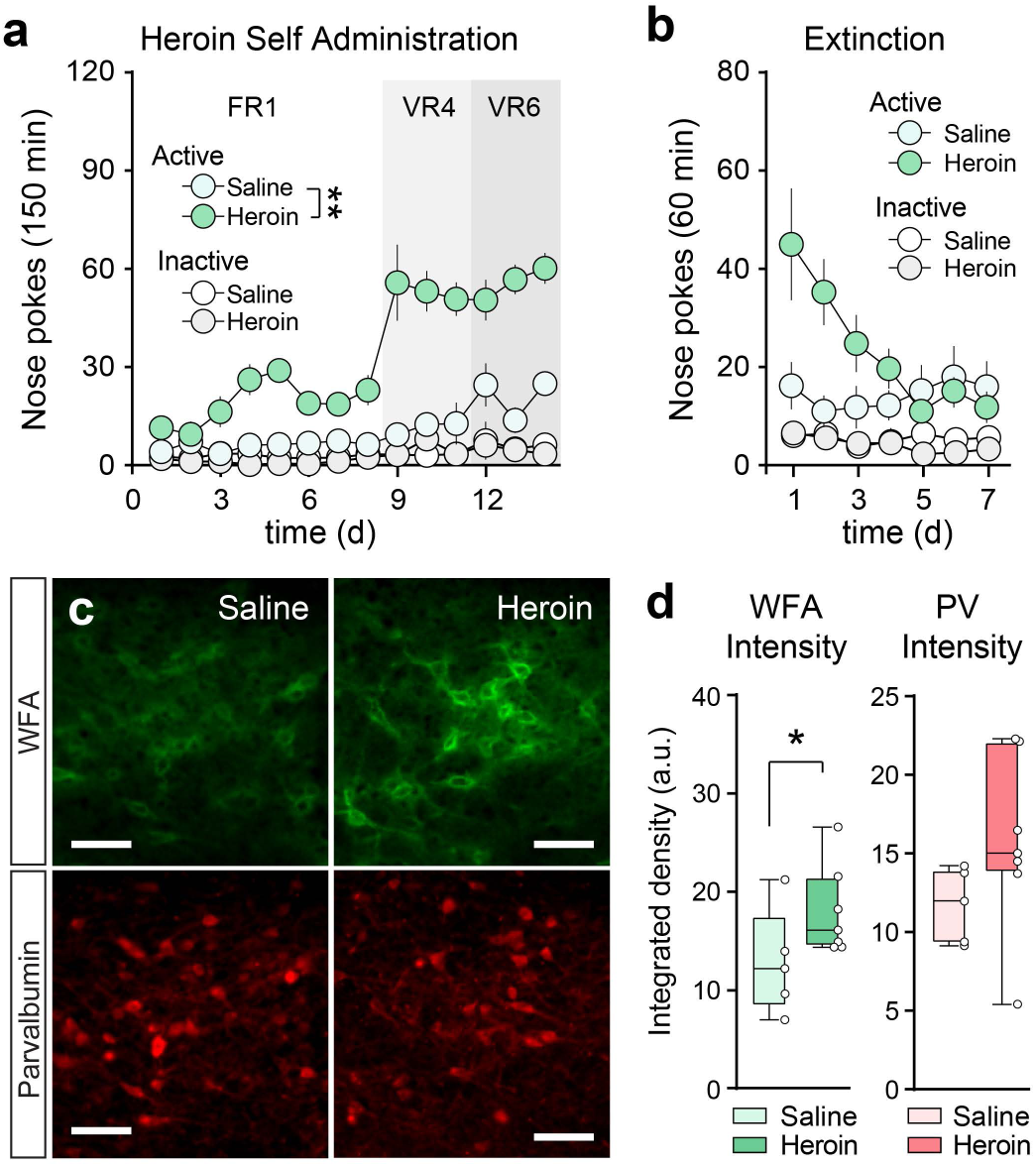
Heroin self-administration increases PNN density in the VP. (**a**) Intravenous heroin and saline self-administration and (**b**) extinction in mice. Data (a,b) are shown as mean ± S.E.M. (**c**) Representative micrographs showing WFA (green) and PV labeling (red) from a heroin and saline self-administering mouse. Scale bar = 100 μm (**d**) Heroin self-administration and withdrawal (extinction) increases WFA intensity without altering PV density. WFA intensity (integrated density; arbitrary units) was shifted by -35 units and PV intensity was shifted by -50 units for graphing purposes. Data (d) are shown as median with interquartile range, and symbols represent individual mice (n = 5-7 per group). * p < 0.05, ** p < 0.01 comparing between saline and heroin.

### Ablating PNNs in the VP reduces heroin seeking

To determine whether VP PNNs are necessary for relapse to heroin seeking (cue-induced reinstatement), we next implanted a group of mice with bilateral cannulae targeting the VP and intravenous jugular vein catheters to examine the effects of PNN ablation on cue-induced heroin seeking. After 14d of heroin self-administration and 7d of extinction training mice received an intracranial microinfusion of ChABC or 0.1% bovine serum albumin containing phosphate buffered saline (PBS) vehicle in their home cage (i.e. on the 8^th^ day of abstinence from heroin) and were subjected to a cue-induced reinstatement test the next day (see: timeline in Figure 3a). Microinfusions of ChABC, but not PBS, significantly reduced the number of PNNs in the VP one day after treatment, as verified by immunostaining for PV and substance P, and WFA-lectin PNN labeling (Figure 3b-c; unpaired t-test: t_(12)_ = 13.29, p = 1.538 × 10^-8^). Cannulated mice rapidly acquired heroin self-administration and extinguished their responding in the absence of heroin and conditioned cues prior to reinstatement testing (Figure 3e-f). Ablating VP PNNs with ChABC significantly reduced cue-induced reinstatement of heroin seeking, measured by active nose pokes (Figure 3g: two-way RM-ANOVA significant main effects of reinstatement: F_(1, 15)_ = 20.33, p = 4.159 × 10^-4^, and treatment x reinstatement interaction: F_(1, 15)_ = 5.837, p = 0.029). Post-hoc tests revealed significant reinstatement in the PBS vehicle control group compared to extinction baseline, but no reinstatement in the ChABC group. No effects were observed on inactive responses (Figure 3h).

**Figure 3.**
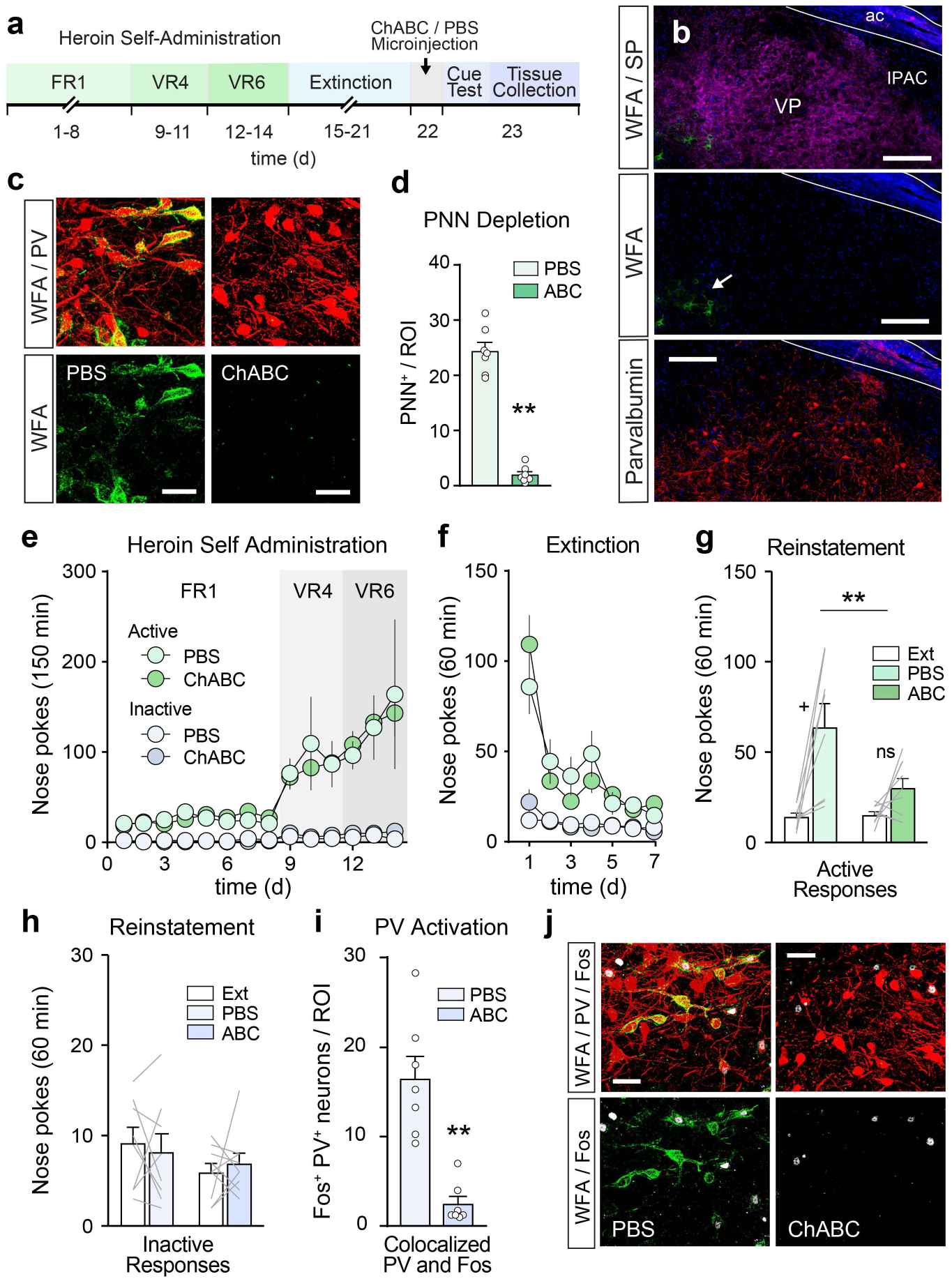
Depletion of VP PNNs reduces heroin seeking. (**a**) Experimental timeline for PNN depletion experiment. Mice underwent 14d of heroin self-administration followed by 7d of extinction training. Afterwards mice received a microinfusion of ChABC or PBS vehicle in their home cage, followed one day later by cue-induced reinstatement testing and tissue collection for Fos immunohistochemistry. (**b**) Representative image showing that ChABC depletes PNNs across the VP. ac = anterior commissure, IPAC = interstitial nucleus of the posterior limb of the anterior commissure. Scale bar = 150 μm. Arrow indicates the edge of the PNN depleted area. (**c**) High magnification micrographs showing the depletion of PNNs around VP_PV_ neurons following ChABC treatment. Scale bar = 40 μm. (**d**) ChABC reduces the number of PNNs in the VP. (**e-f**) Cannulated mice acquired heroin self-administration and extinguished their responding in the absence of heroin and conditioned stimuli. (**g**) ChABC depletion of PNNs in the VP reduced cue-induced reinstatement of heroin seeking, measured by active nose pokes. (**h**) PNN depletion did not alter inactive nose pokes. (**i**) PNN depletion significantly reduce the number of Fos positive VP_PV_ neurons during cue-induced reinstatement. (**j**) Representative micrograph showing a lack of Fos (white) in VP_PV_ (red) neurons during reinstatement following depletion of PNNs (green). Scale bar = 40 μm. ABC = chondroitinase ABC, PBS = phosphate buffered saline vehicle. Symbols in bars (dots or lines) represent individual mice per treatment group. ** p < 0.01 comparing treatment groups. + p < 0.05 comparing between extinction baseline and cue-induced reinstatement. Data are shown as mean ± s.e.m.

To examine the effects of PNN depletion on the activation of VP_PV_ neurons during cue-induced reinstatement, we transcardially perfused mice one hour after completion of the reinstatement test and immunostained their brains for the immediate early gene product Fos and PV to investigate the activation of VP_PV_ neurons. Cell counts showed a significantly reduced activation of VP_PV_ neurons during reinstatement in the ChABC group compared to PBS vehicle (Figure 3i-j: unpaired t-test: t_(12)_ = 5.275, p = 1.964 × 10^-4^), indicating a reduced activation of VP_PV_ neurons during reinstatement following PNN depletion. Only mice with confirmed microinjection placements in the VP were included in analyses (Supplementary Figure 1). Overall, these results show that PNN depletion reduces heroin seeking and Fos expression in VP_PV_ neurons and suggest that VP PNNs regulate the activity of VP_PV_ neurons to drive relapse to heroin seeking.

### Ablating VP PNNs alters the intrinsic excitability, membrane properties and synaptic inputs of VP_PV_ neurons

Next, we examined the effects of ChABC microinfusions in the VP on the intrinsic excitability of VP_PV_ neurons and their synaptic inputs using ex-vivo patch-clamp electrophysiology. PV-tdTomato reporter mice received a randomized infusion of ChABC in one hemisphere and a contralateral infusion of PBS vehicle. Brains for electrophysiological analyses were collected 24h later. The localization of recorded cells within the PNN depleted area of the VP was verified post-recording using a combination of biotin labeling, immunohistochemistry for tdTomato, and WFA-lectin staining (Figure 4a; Supplementary Figure 2a). Depleting PNNs significantly reduced the intrinsic excitability of VP_PV_ neurons and reduced their maximum firing rates across a broad range of current injection steps (Figure 4b,c; Two-way RM-ANOVA main effect of current step: F_(20, 500)_ = 1.110 × 10^-16^, treatment: F_(1,25)_ = 15.28, p = 6.260 × 10^-4^, and interaction: F_(20, 500)_ = 2.134, p = 0.003).

**Figure 4.**
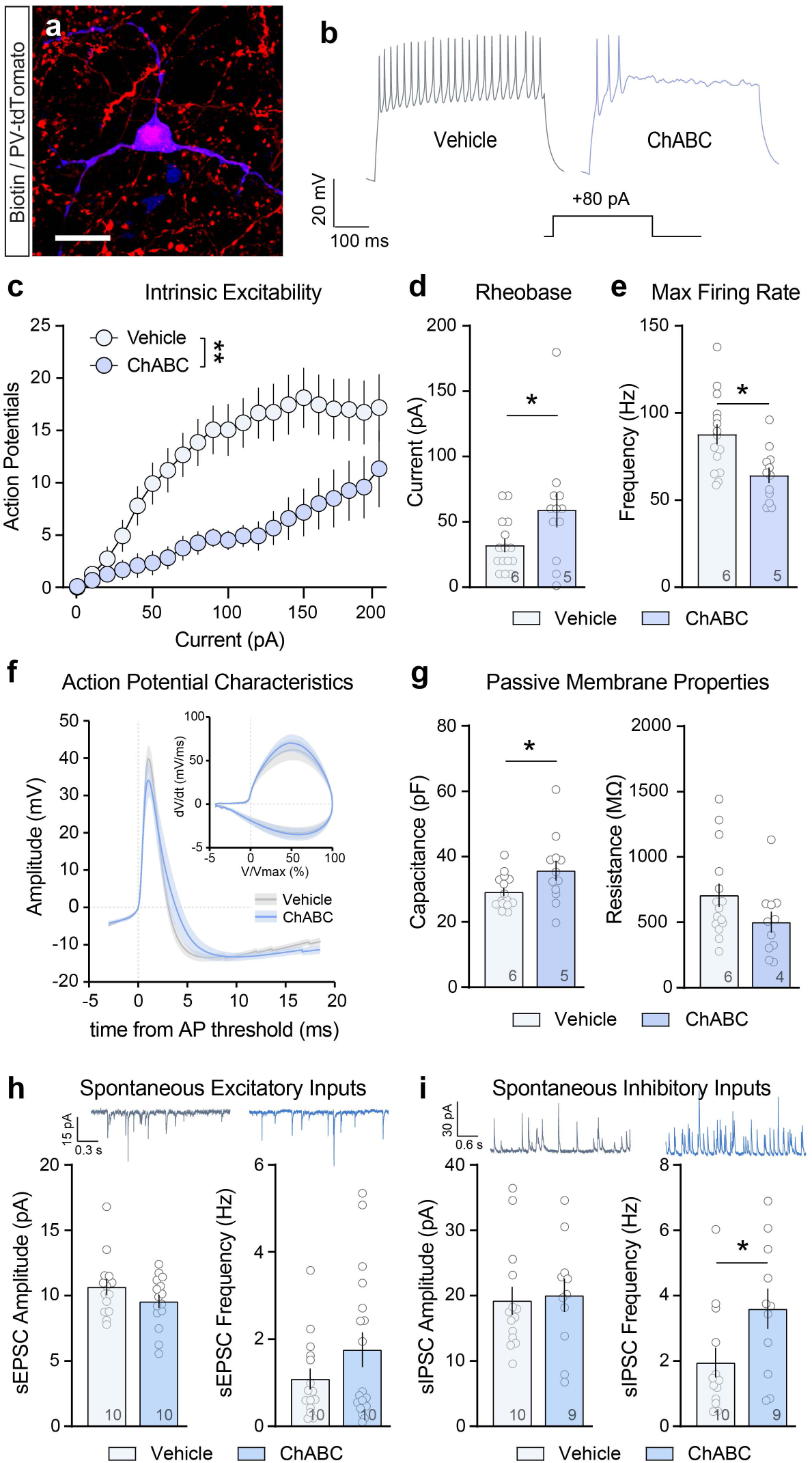
VP PNN depletion reduces the intrinsic excitability and membrane properties of VP_PV_ neurons and alters synaptic inputs onto these neurons. (**a**) Representative image showing a biotin (blue) filled VP_PV_ (red) neuron recorded from a PV-tdTomato mouse. Scale bar = 20 μm. (**b**) Representative traces showing that ChABC-mediated PNN depletion (blue) reduces the number of action potentials recorded from a VP_PV_ neuron following an 80 pA current injection step. (**c**) PNN depletion reduces the intrinsic excitability of VP_PV_ neurons measured at different current injection steps. (**d**) ChABC treatment increases the rheobase of VP_PV_ neurons, and (**e**) reduces the maximum firing rate. (**f**) PNN depletion does not alter the average shape or phase-plane representation of the action potential. (**g**) ChABC treatment increases the membrane capacitance, without altering membrane resistance. (**h)** PNN depletion does not alter sEPSC frequency or amplitude, but (**i**) increases sIPSC frequency onto VP_PV_ neurons, without altering sIPSC amplitude. Representative traces show sEPSC and sIPSC recorded from VP_PV_ neurons after PBS (grey) or ChABC treatment (blue). Symbols in bars represent the number of recorded cells and numbers in bars represent the number of mice. ** p < 0.01, * p < 0.05 comparing between vehicle and ChABC treated cells. Data shown as mean ± s.e.m.

Similarly, ChABC increased the rheobase of VP_PV_ neurons (Figure 4d; Unpaired t-test: t_(25)_ = 2.076, p = 0.048) and reduced their maximum firing rate (Figure 4e; Unpaired t-test: t_(25)_ = 3.111, p = 0.005). Although PNN depletion did not alter the average shape of VP_PV_ action potentials (Figure 4f, Supplementary Figure 2c-h), it did increase the number of neurons with a two-component action potential rising phase in the phase-plane plot (Supplementary Figure 2i-k; Chi-square test: χ^2^_(1)_ = 4.531, p = 0.033). VP PNN depletion also altered the passive membrane properties of VP_PV_ neurons, significantly increasing membrane capacitance following ChABC treatment (Figure 4g; Unpaired t-test: t_(25)_ = 2.114, p = 0.045) without changing membrane resistance. In addition, we examined synaptic changes onto VP_PV_ neurons following ChABC treatment and found that depleting PNNs increased the frequency of spontaneous inhibitory post synaptic currents (sIPSC; Figure 4h-i; Supplementary Figure 3a-b; Unpaired t-test: t(22) = 2.197, p = 0.039) without altering sIPSC amplitude or spontaneous excitatory postsynaptic current (sEPSC) amplitude or frequency. Neither evoked nor spontaneous excitation/inhibition ratios, nor paired pulse ratios were changed following PNN depletion (Supplementary Figure 3c-h). Combined, these data suggest that PNN depletion reduces the excitability of VP_PV_ neurons through an increase in membrane capacitance, and a concurrent increase in spontaneous inhibitory synaptic inputs.

### Chemogenetic stimulation of VP_PV_ neurons rescues heroin seeking following ChABC treatment

Given that PNN depletion reduces Fos and the intrinsic excitability of VP_PV_ neurons, we next hypothesized that chemogenetic activation of these neurons would be sufficient to rescue heroin seeking in ChABC-treated mice. PV-IRES-Cre mice were injected with AAV-hsyn-DIO-hM3D-mCherry in the VP to express the Gq-coupled excitatory designer receptor exclusive activated by designer drugs (Gq DREADD) hM3D in VP_PV_ neurons. During the same surgery, mice were implanted with chronic guide cannulas for ChABC delivery, and jugular vein catheters. Virus expression and cannula placements were confirmed using immunohistochemistry for mCherry and substance P, and WFA-lectin labeling for PNNs (Figure 5a, b, Supplementary Figure 4a). Cannula-implanted and AAV injected mice readily acquired heroin self-administration over 14d and extinguished their responding over a subsequent 7d of extinction training. (Figure 5c, d). 24 hours prior to reinstatement testing mice received an intracranial microinfusion of ChABC or PBS vehicle and the next day they received an intraperitoneal (i.p.) injection of saline vehicle or the chemogenetic ligand J60 (Bonaventura et al., 2019), 30 minutes prior to testing. Ex-vivo patch-clamp electrophysiological validation of the viral vector functionally confirmed that bath-application of J60 significantly increased the firing of VP_PV_ neurons (Supplementary Figure 4b; Wilcoxon signed-rank test: n = 10, W = 50, p = 0.008), and that it raised the membrane potential of recorded neurons (Wilcoxon signed-rank test: n = 13, W = 91, p = 2.441 × 10^-4^). Consistent with our previous findings, ChABC microinjections in the VP significantly reduced active lever responding for heroin in saline treated mice, compared to PBS microinjected and saline treated control mice, and chemogenetic activation of VP_PV_ neurons with J60 reversed this effect (Figure 5e – left; active nose poke two-way RM-ANOVA main effect of reinstatement: F_(1,20)_ = 41.85, p = 2.623 × 10^-6^, and treatment x reinstatement interaction F_(2,20)_ = 5.573, p = 0.012). Post-hoc tests confirmed that ChABC-treated mice failed to reinstate their heroin seeking, and that chemogenetic activation of VP_PV_ neurons successfully rescued heroin seeking to the level seen in PBS microinjected controls. No effects were observed on inactive nose pokes (Figure 5e – right). These findings support the notion that VP PNN depletion reduces heroin seeking by reducing the activity of VP_PV_ neurons.

**Figure 5.**
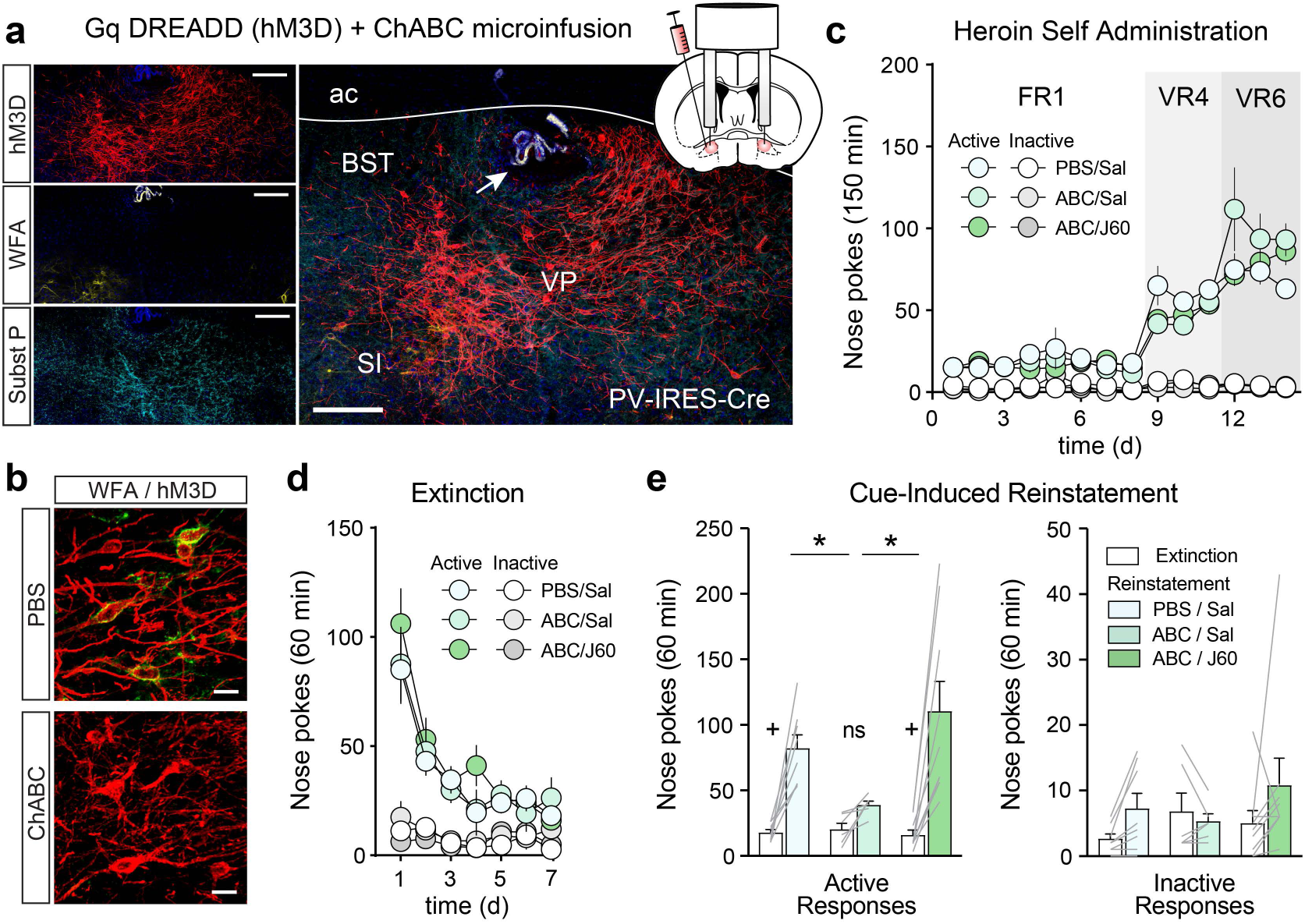
Chemogenetic stimulation of VP_PV_ neurons prevents the effects of ChABC on heroin seeking. (**a**) Representative image of Gq DREADD expression in VP_PV_ neurons of a PV-IRES-Cre mouse (red), PNN depletion after ChABC microinfusion (yellow), and substance P staining to delineate the boundaries of the VP (teal). Arrow indicates cannula placement in the VP. Scale bar = 150 μm. ac = anterior commissure, BST = bed nucleus of the stria terminalis, SI = substantia innominata. (**b**) High magnification micrograph showing ChABC mediated depletion of PNNs around virally transduced VP_PV_ neurons. Scale bar = 20 μm. (**c**) Heroin self-administration and (**d**) extinction in mice implanted with cannulas and injected with AAV-hSyn-DIO-hM3D-mCherry. (**e**) Chemogenetic stimulation of VP_PV_ neurons with the hM3D ligand J60 reverses the effects of PNN depletion on heroin reinstatement. Lines in bars represent individual mice. * p < 0.05 comparing active nose pokes during reinstatement between groups. + p < 0.001 comparing between extinction baseline and reinstatement for each group. Data shown as mean ± s.e.m.

### Chemogenetic inhibition of VP_PV_ neurons mimics the effects of ChABC on heroin seeking

Since ablating PNNs induced a hypoactive state in VP_PV_ neurons that suppressed heroin seeking, we hypothesized that direct chemogenetic inhibition of VP_PV_ neurons would recapitulate this behavioral effect. PV-IRES-Cre mice were injected with the inhibitory Gi-coupled DREADD hM4D construct AAV-hSyn-DIO-hM4D-mCherry or an AAV-hSyn-DIO-mCherry control virus in the VP (Figure 6a). Functional ex-vivo validation experiments confirmed that J60 significantly reduced the firing rate of VP_PV_ neurons (Supplementary Figure S6c; Wilcoxon signed-rank test: n = 10, W = -47, p = 0.014) and hyperpolarized the membrane potential (Wilcoxon signed-rank test: n = 10, W = -55, p = 0.002) of hM4D transduced VP_PV_ neurons. Mice expressing hM4D or mCherry in VP_PV_ neurons readily acquired heroin self-administration over 14d and extinguished their responding over 7d of extinction training (Figure 6b,c). Afterwards, mice received an i.p. injection of J60 or saline vehicle, 30 minutes prior to testing. Chemogenetic inhibition of VP_PV_ neurons significantly attenuated reinstatement of heroin seeking compared to hM4D-expressing saline controls or J60-treated mCherry-expressing mice (Figure 6d; active nose poke two-way RM-ANOVA main effect of reinstatement: F_(1, 18)_ = 38.78, p = 7.098 × 10^-6^, and treatment x reinstatement interaction: F_(2, 18)_ = 7.660, p = 0.004). Post-hoc tests revealed that while the control groups exhibited robust reinstatement, hM4D-expressing mice treated with J60 failed to reinstate heroin seeking. No effects were observed on inactive nose pokes. These results demonstrate that silencing VP_PV_ neurons is sufficient to recapitulate the anti-relapse effects of VP PNN depletion.

**Figure 6.**
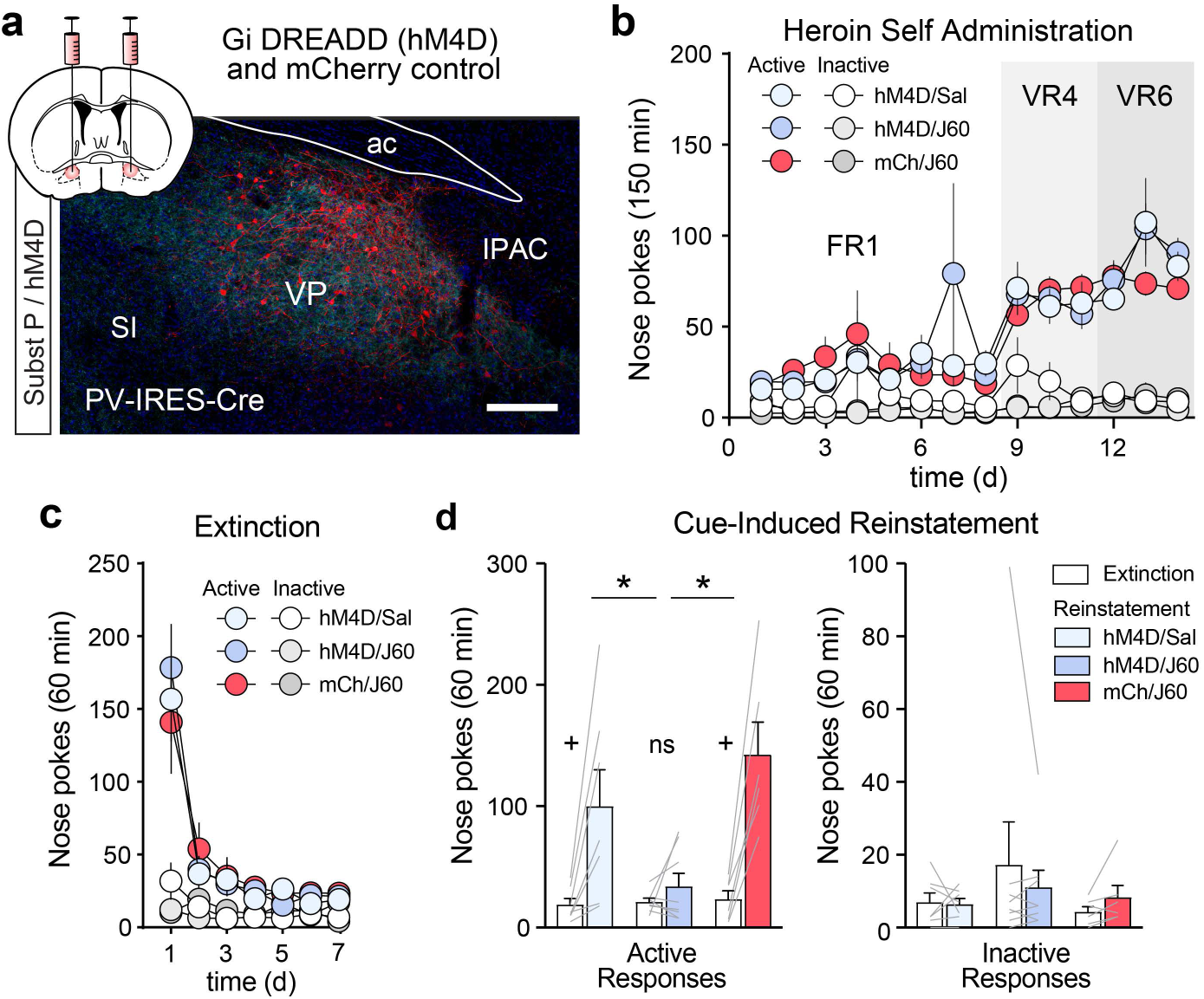
Chemogenetic inhibition of VP_PV_ neurons mimics the effects of ChABC on heroin seeking. (**a**) Representative micrograph showing hM4D-mCherry expression in VP_PV_ neurons (red) within the boundaries of the VP as determined by a substance P counterstain (teal). ac = anterior commissure, SI = substantia innomunata, IPAC = interstitial nucleus of the posterior limb of the anterior commissure. Scale bar = 500 μm. (**b**) Heroin self-administration and (**c**) extinction responding in mice transduced with AAV-hSyn-DIO-hM4D-mCherry or AAV-hSyn-DIO-mCherry. (**d**) Chemogenetic inhibition of VP_PV_ neurons with the hM4D ligand J60 reduced heroin reinstatement. No effects were observed on inactive nose pokes. Lines in bars represent individual mice. * p < 0.05 comparing active nose pokes during reinstatement between groups, + p < 0.01 comparing between extinction and reinstatement for each group. Data shown as mean ± s.e.m.

## Discussion

In the present study, we identified VP PNNs as a key structural regulator of cued relapse to opioid seeking. By combining chemogenetics with the enzymatic depletion of PNNs, we demonstrate that the structural integrity of PNNs in the VP is necessary for the activation of VP_PV_ neurons and heroin seeking, and that stimulating VP_PV_ neurons is sufficient to rescue heroin seeking after VP PNN depletion. Our work also corroborates prior reports showing that the VP contains a high number of PNNs and firmly establishes that PNNs preferentially localize to VP_PV_ neurons (Brauer et al., 1993; Seeger et al., 1994). In addition, our findings that heroin increases the density of VP PNNs and that their removal from the VP reduces opioid seeking add to a growing literature linking structural changes in PNNs to motivation for drugs of abuse (Fawcett et al., 2022; Lubbers et al., 2014). Taken together, our results show that the functional regulation of VP_PV_ neurons by PNNs is critical for relapse to heroin seeking.

While prior work indicated limited to no co-localization of PNNs to VP_PV_ neurons (Brauer et al., 1993), our results indicate that a majority of VP_PV_ neurons are surrounded by PNNs. This discrepancy is likely explained by methodological differences in immunohistochemical methods or WFA-lectin staining between our studies. Nonetheless, we did observe a minor subpopulation of PNN positive cells in the VP that did not appear to express PV. Similarly, a prior study reported that a subset of PNN positive neurons in the striatum does not express PV (Lee et al., 2012). It is possible that these neurons express levels of PV below the immunohistochemical detection threshold, and that more sensitive methods may uncover low PV expression. Alternatively, these neurons may represent a distinct, non-parvalbumin expressing population surrounded by PNNs, as is for instance observed in the CA2 region of the hippocampus (Carstens et al., 2016).

Our ex-vivo patch-clamp electrophysiology analyses revealed a reduced intrinsic excitability of VP_PV_ neurons following PNN depletion. Several mechanisms could mediate this effect. Consistent with prior studies, our data show that PNN depletion increased membrane capacitance, thereby increasing the effective membrane surface area (Tewari et al., 2018; Wingert et al., 2024). Neurons with a higher membrane capacitance require more current to reach action potential threshold, which likely contributed directly to our observed increase in rheobase and reduced firing rates. In addition, PNNs are known to constrain the mobility of voltage gated ion-channels and their negative charge may facilitate cation buffering near the soma to regulate intrinsic excitability (Hartig et al., 1999; Morawski et al., 2015; Wingert & Sorg, 2021). Although most studies report a decrease in the excitability of PV neurons following PNN depletion (Wingert et al., 2024; Wingert & Sorg, 2021), some studies suggest a subsequent compensatory increase in excitability (Slaker et al., 2015). Importantly, the effects of ChABC are transient, as PNNs gradually regenerate around PV neurons over the course of weeks, and this recovery is thought to once again terminate a window of heightened plasticity within the targeted brain region (Slaker et al., 2015). Future work should examine the precise temporal dynamics of this structural remodeling in the VP and its potential effects on subsequent heroin seeking.

Diverse synaptic changes have been reported in PV neurons across different brain regions following PNN depletion (Wingert & Sorg, 2021). We observed a specific increase in the frequency of spontaneous inhibitory postsynaptic currents (sIPSCs) onto VP_PV_ neurons. PNNs are known to constrain the diffusion of ligand-gated ion channels, which may have contributed to this effect (Frischknecht et al., 2009; Wingert & Sorg, 2021). Additionally, PNN depletion facilitates the formation of new synaptic contacts onto PV neurons (Slaker et al., 2018). A specific increase in the frequency of inhibitory postsynaptic inputs onto PV neurons was previously reported in the hippocampus following PNN depletion (Hayani et al., 2018). However, any changes to synaptic inputs following PNN depletion are likely highly specific to the local circuitry of the studied brain region. VP_PV_ neurons do receive very dense inhibitory inputs from the nucleus accumbens, and GABAergic synapses in the VP outnumber glutamatergic synapses at least 5-fold (Chang et al., 1995; Do et al., 2016; Knowland et al., 2017). Thus, a potential explanation for a selective increase in the sIPSC frequency onto VP_PV_ neurons that we observed could be that removing VP PNNs specifically unmasked this dense inhibitory tone in the VP.

Our data show that PNN depletion does not alter the action potential characteristics of VP_PV_ neurons, whereas other studies report a slowing of the action potential half width and an increased delay to afterhyperpolarization (Wingert & Sorg, 2021). In part, this discrepancy could be explained by a high degree of variability in the shape of action potentials that we recorded from VP_PV_ neurons (i.e. Supplementary Figure 4i,j). Irrespective of this variability, we observed a significant shift in the number of VP_PV_ neurons with a two-component rising phase of the action potential in the phase-plane plot after PNN depletion. Such a change is thought to be mediated by reduced resistive coupling between the axon initial segment (AIS) and the somatodendritic compartment of a neuron, and likely mediated by the increased capacitance of VP_PV_ neurons that we observed after PNN depletion (Bean, 2007; Kole & Brette, 2018). Alternatively, changes in the location and biophysical properties of the AIS have been proposed to contribute to changes in the firing properties of a neuron and the resistive coupling between cellular compartments (Caspi et al., 2023; Kole & Brette, 2018). Since most PNNs cover the AIS, PNN depletion at this specific locus may have played a key role in the observed changes in neuronal excitability following PNN depletion (Bruckner et al., 2006; Caspi et al., 2023).

Congruent with a previous study on alcohol seeking in rats (Prasad et al., 2020), we show that VP_PV_ neurons regulate cue-induced opioid seeking in mice. Given that the activation VP_PV_ neurons is required for drug seeking behavior, and that PNNs increase the intrinsic excitability of PV neurons, the increased density of PNNs surrounding VP_PV_ neurons after heroin self-administration that we observed likely contributed to heroin seeking by regulating the activity of VP_PV_ neurons. Indeed, our chemogenetic experiments provide a causal link between physiological changes caused by PNN depletion and observed reductions in reinstatement. We found that chemogenetic inhibition of VP_PV_ neurons recapitulated the effects of VP PNN depletion, confirming that the activity in this population of neurons is necessary for heroin seeking. Crucially, our findings also showed that chemogenetic stimulation of VP_PV_ neurons was sufficient to rescue heroin seeking following PNN depletion by ChABC. Collectively, these findings indicate that an increased density of VP PNNs after heroin self-administration acts to facilitate the activation of VP_PV_ neurons to drive heroin seeking. However, it is important to note that VP_PV_ neurons form a heterogenous population with different cellular properties and behavioral functions depending on the neurotransmitters they produce and their projection targets (Knowland et al., 2017; Prasad et al., 2020; Tooley et al., 2018).

The VP contains both glutamatergic and GABAergic VP_PV_ neurons (Graham et al., 2024; Knowland et al., 2017; Tooley et al., 2018) which have distinct functions in reward processing (Soares-Cunha & Heinsbroek, 2023), and VP GABAergic (VP_GABA_) and glutamatergic (VP_Glu_) neurons exert opposite control over motivated behavior (Faget et al., 2018; Farrell et al., 2021; Heinsbroek et al., 2020; Stephenson-Jones et al., 2020; Tooley et al., 2018). Since VP_GABA_ neurons are established drivers of relapse to heroin and cocaine seeking (Farrell et al., 2022; Heinsbroek et al., 2020), and both VP_GABA_ and VP_PV_ neurons drive relapse to alcohol seeking (Prasad et al., 2020), it is likely that a GABAergic subpopulation of VP_PV_ neurons was responsible for driving heroin seeking in our study. Nonetheless, a role for glutamatergic VP_PV_ neurons cannot be excluded at this time given their known role in aversive states associated with drug withdrawal, arousal, and salience processing, which may each contribute to drug seeking (Levi et al., 2025; Wang et al., 2020; Xu et al., 2025). Taken together, VP PNNs likely play a nuanced role in the regulation opioid seeking and future studies should employ cell- and circuit specific PNN depletion strategies (Favuzzi et al., 2017; Hazlett et al., 2024) to parse these distinct circuit mechanisms.

PNNs have been hypothesized to regulate the long-term stability of neural circuits, and to facilitate memory storage (Tsien, 2013). However, rapid and transient changes in PNN structure have also been reported (Fawcett et al., 2022; Harkness et al., 2021). Notably, the extracellular matrix has been shown to undergo rapid changes in the nucleus accumbens during reinstatement, which is associated with a heightened motivation to seek drugs of abuse (Smith et al., 2014; Smith et al., 2017). Whether VP PNNs are capable of similar rapid structural changes remains an important open question that can provide key additional insights into the VP processes that govern relapse to opioid seeking.

In conclusion, our work establishes that VP PNNs and the VP_PV_ neurons they surround are critical regulators of opioid seeking. These findings set the stage for additional investigation into cell- and circuit-specific molecular, structural and functional adaptations in VP PNNs induced by opioids, and their potential contributions to persistent craving and relapse risk in opioid use disorder.

## STAR * METHODS

### Key resources table

| REAGENT or RESOURCE | SOURCE | IDENTIFIER |
| --- | --- | --- |
| **Antibodies** |  |  |
| Chicken anti-mCherry (1:10,000) | Synaptic Systems | LS-C204825;<br>RRID:AB_2716246 |
| Guinea Pig anti-RFP (1:1,000) | Synaptic Systems | Cat#390 004;<br>RRID:AB_2737052 |
| Parvalbumin (1:1,000) | Synaptic Systems | Cat#195 004;<br>RRID:AB_2156476 |
| Rabbit anti-dsRed (1:1,000) | Clontech | Cat#632496;<br>RRID:AB_10013483 |
| Substance P (1:1,000) | Immunostar | Cat#20064,<br>RRID:AB_572266 |
| Phospho-Ser32 Fos (1:5,000) | Cell Signaling<br>Technology | Cat#53485,<br>RRID:AB_10557109 |
| Wisteria Floribunda Agglutinin (WFA), fluorescein-<br>conjugated (1:500) | Vector Laboratories | Cat#: FL-1351;<br>RRID:AB_2336875 |
| Wisteria Floribunda Agglutinin (WFA), biotin-conjugated<br>(1:200) | Vector Laboratories | Cat#: B-1355;<br>RRID:AB_2336874 |
| Glycoprotein-Free Blocking Solution | Vector Laboratories | Cat#: SP-5040-125 |
| DAPI-containing mounting medium (ProLong™<br>Diamond) | Invitrogen / Thermo<br>Fisher Scientific | Cat#: P36962 |
| **Bacterial and virus strains** |  |  |
| AAV2-hSyn-DIO-hM3D(Gq)-mCherry | Addgene | Cat# 44361-AAV2,<br>RRID: Addgene_44361 |
| AAV2-hSyn-DIO-hM4D(Gi)-mCherry | Addgene | Cat# 44362-AAV2,<br>RRID: Addgene_44362 |
| AAV2-hSyn-DIO-mCherry | Addgene | Cat# 50459-AAV2,<br>RRID: Addgene_50459 |
| **Chemicals, peptides, and recombinant proteins** |  |  |
| Diamorphine hydrochloride (Heroin) | NIDA Drug Supply<br>Program | Cat#: 9200-001<br>CAS: 1502-95-0 |
| Chondroitinase ABC (ChABC) | Sigma-Aldrich | Cat#: C3667-5UN;<br>CAS: 9024-13-9 |
| Bovine serum albumin (BSA) | Sigma-Aldrich | Cat#: A8806;<br>CAS: 9048-46-8 |
| Phosphate buffered saline (PBS) | Thermo Fisher<br>Scientific | Cat#: 10010023;<br>CAS: 7732-18-5 |
| JHU37160 dihydrochloride (J60) | Hello Bio | Cat# HB6261<br>CAS: 2369979-68-8 |
| Isoflurane | Sigma-Aldrich | Cat#: 792632;<br>CAS: 26675-46-7 |
| Carprofen (Rimadyl) | Zoetis | Cat#: 50-282-8028<br>CAS: 53716-49-7 |
| Cefazolin | Hikma | Cat#: 00143992490<br>CAS: 5331-34-6 |
| Taurolidine-citrate solution (TCS) | Access Technologies | Cat#: NC0444187<br>CAS: 1333382-80-1 |
| Heparin (50,000 USP, 10 ml stock solution) | Fresenius Kabi LLC | Cat#: 4710<br>CAS: 9005-49-6 |
| Sodium Methohexital | PAR Pharmaceuticals | Cat#: 42023010501; CAS:<br>309-36-4 |
| Euthasol® | Virbac AH, Inc. | Cat#: 51311-050-01<br>CAS: 57-33-0 |
| Formalin (10%) | Fisher Bioreagents | Cat#: F/1565/53;<br>CAS: 50-00-0 |
| Isopentane | Sigma-Aldrich | Cat#: 277258;<br>CAS: 78-78-4 |
| NBQX disodium salt hydrate | Sigma-Aldrich | Cat# N183;<br>CAS: 479347-86-9 |
| Picrotoxin | Sigma-Aldrich | Cat# P1675;<br>CAS: 124-87-8 |
| **Deposited data** |  |  |
| Data | This paper |  |
| **Experimental models: Organisms/strains** |  |  |
| Mouse: PV-tdTomato | The Jackson Laboratory | RRID: IMSR_JAX:027395 |
| Mouse: PV-IRES-Cre | The Jackson Laboratory | RRID: IMSR_JAX:017320 |
| Mouse: C57BL/6J | The Jackson Laboratory | RRID:IMSR_JAX:000664 |
| **Software and algorithms** |  |  |
| Fiji | ImageJ; NIH | RRID:SCR_002285 |
| pClamp v11 | Molecular Devices | RRID:SCR_011323 |
| ClampEx v11 | Molecular Devices | RRID:SCR_011323 |
| Clampfit v11 | Molecular Devices | RRID:SCR_011323 |
| MATLAB R2023b | Mathworks | RRID:SCR_001622 |
| GraphPad Prism v9.0.0 | GraphPad | RRID:SCR_002798 |

### Subjects

Male and female mice were used for all procedures. Mice were kept on a regular 12h day/night cycle and were group housed and provided with food and water ad libitum until the start of experiments. PV-tdTomato (Jackson labs: #027395), PV-IRES-Cre (#017320) and wildtype mice were bred in house on a C57BL/6J background. All experiments were approved by the institutional animal care and use committees at the University of Colorado and the University of Alabama at Birmingham.

### Drugs

Diamorphine hydrochloride (Heroin) was generously provided by the National Institute on Drug Abuse and Research Triangle Institute, dissolved in sterile saline and filtered. Heroin was self-administered at 150 μg/kg in a volume of 12 μl per infusion. Chondroitinase ABC (ChABC) was obtained from Sigma (C3667-5UN) and dissolved in sterile 0.1% bovine serum albumin (BSA) containing phosphate buffered saline (PBS) to an effective concentration of 50 U/ml. 24 h prior to reinstatement testing ChABC was microinfused into the VP in a volume of 100 nl at a rate of 50 nl per min using microinjectors that extended 1 mm below the tip of the cannula. Following the infusion, microinjectors were left in place for an additional 4 min to allow for the diffusion of ChABC. JHU37160 dihydrochloride (J60; HB6261) was obtained from Hello Bio, dissolved in sterile saline and administered intraperitoneally (i.p.) at 1 mg/kg (10 ml/kg), 30 minutes prior to testing.

### Virus

All adeno-associated viral (AAV) vectors were obtained from Addgene. AAV2-hSyn-DIO-hM3D(Gq)-mCherry, AAV2-hSyn-DIO-hM4D(Gi)-mCherry, and AAV2-hSyn-DIO-mCherry were diluted to 5 × 10^12^ GC/ml in sterile PBS and infused bilaterally at a volume of 150 nl using a nanoliter pressure injection system (15 injections of 10 nl at a rate of 1 nl/s, with a 20s interval between injections). Virus was given at least 10 min to diffuse, after which glass pipettes were slowly retracted.

### Surgery

Mice were anesthetized with isoflurane (induction: 5%, maintenance: 1-2% v/v) and were given carprofen (5 mg/kg; i.p.) for analgesia and cefazolin (100 mg/kg) to prevent infection. For catheter surgeries an indwelling catheter (Instech) was implanted 11 mm into the jugular vein and connected to an access port between the scapulae on the back. To maintain patency, catheters were filled with taurolidine-citrate solution (Access Technologies) during surgery and flushed daily with cefazolin and sterile heparinized saline (100 IU/ml) during self-administration. Catheter patency was tested using sodium methohexital (5 mg/ml), and mice with catheters that had lost patency were excluded from experiments. For cranial surgery, mice were mounted in a stereotaxic frame (Kopf). Bilateral intracranial cannulas (P1 technologies) were slowly lowered above the VP using the following coordinates: anteroposterior (AP), mediolateral (ML) and dorsoventral (DV) relative to Bregma; AP: 0.4, ML: ±1.5, DV: -4.1. Cannulas were anchored to the skull using jewelers’ screws and dental cement, and dummy injectors were inserted to prevent obstruction. Virus was infused using the following coordinates: AP: 0.4, ML: ±2.4 (10° angle), DV: -4.8. After surgery mice were given at least 1w to recover, and testing was conducted at least 3w after virus infusion to allow for sufficient expression. Following surgery mice were single housed throughout all experiments with limited environmental enrichment, as enrichment is a known regulator of PNN structure (Foscarin et al., 2011).

### Behavior

Mice were habituated to handling and catheter flushing for at least 3d prior to starting heroin self-administration. Afterwards, mice started daily 150 min heroin self-administration sessions on a fixed ratio (FR1) schedule of reinforcement in modular operant chambers (Med Associates). A house light and fan were on during the entirety of the session, and a cue light over top of the ‘active’ nosepoke hole signaled heroin availability. Active nose pokes that yielded heroin infusions led to the presentation of a 2s compound cue (2 kHz tone and the illumination of the noke poke port). The availability light was off during reward delivery and during a subsequent 10s time-out period. After 8 days, mice progressed to a variable ratio schedule of reinforcement (VR4), followed by VR6. Then, mice underwent 1h extinction training sessions for 7d, followed by a day of abstinence in the home cage during which VP microinfusions were given.

### Histology and Microscopy

Mice used in Fos analyses were perfused 2h after the start of the reinstatement session. In brief, mice were deeply anesthetized with pentobarbital (Euthasol; 150 mg/kg) and transcardially perfused with 10 ml of a custom exsanguination solution (0.9% NaCl, 1% NaNO_3_, 40 IU/ml heparin) (Marchant et al., 2009), followed by 20 ml formalin (10%). Brains were post-fixed overnight in 10% formalin, cryoprotected in 20% sucrose, flash frozen using isopentane, and sectioned at 35 μm thickness on a cryostat. Tissue was immunostained as described previously (Heinsbroek et al., 2020) using primary antibodies raised in chicken against mCherry (1:10k; LS-C204825; RRID:AB_2716246), raised in Guinea Pig against RFP (1:1000, Synaptic Systems, #390 004; RRID:AB_2737052) and Parvalbumin (1:1000, Synaptic Systems, #195 004; RRID:AB_2156476), and raised in Rabbit against dsRed (1:1000; Clontech #632496; RRID:AB_10013483), substance P (1:1000, Immunostar #20064, RRID:AB_572266), and phospho-Ser32 Fos (1:5000; #53485, Cell Signaling; RRID:AB_10557109). Secondary antibodies raised in goat and/or streptavidin conjugated to Alexa-(1:500), and Alexa Plus (1:1000) dyes (Invitrogen) were used. In brief, sections were washed in PBS on a shaker, blocked using a glycoprotein-free blocking solution (Vector Laboratories; SP-5040-125) containing 0.25% Triton X-100 for 30 minutes at room temperature, incubated in 0.25% PBS-Triton (PBS-T) containing primary antibodies overnight at 4°C, and incubated in PBS-T containing secondary antibodies for 3h at room temperature. Following immunolabeling, perineuronal nets were visualized using Wisteria Floribunda Agglutin (WFA) conjugated to fluorescein (1:500, Vector Laboratories; FL-1351; RRID:AB_2336875) or biotin (1:200, Vector Laboratories; B-1355; RRID:AB_2336874) in PBS-T for 30 minutes at room temperature, followed by a streptavidin labeling step (in PBS-T; 1h at room temperature). Tissue was cover slipped using a DAPI-containing mounting medium (Prolong Diamond). Images were acquired using a confocal microscope (Olympus Fluoview 1200 and Zeiss LSM800) or slide scanning microscope (Keyence BZ-X810) and data were analyzed in Fiji (ImageJ; NIH). Integrated density analyses were performed using the Fiji ROI manager using images from sections taken across the anteroposterior axis of the VP. Values were normalized to the total VP area for each image prior to averaging across all images for each mouse. Cell counting and co-localization analyses were performed using the Fiji Cell Counter plugin and individual values represent average numbers across 4 images stacks for each mouse.

### Electrophysiology

Mice were deeply anesthetized using isoflurane and brains were extracted and sliced in an ice-cold N-methyl-D-gluconate based cutting solution (in mM; 92 NMDG, 2.5 KCl, 1.25 NaH_2_PO_4_, 30 NaHCO_3_, 20 HEPES, 25 D-glucose, 2 thiourea, 5 Na-ascorbate, 3 Na-pyruvate, 0.5 CaCl_2_, and 10 MgSO_4_) (Ting et al., 2018). Brain sections (275 μm) were sliced on a vibratome (Leica VT1200S) and collected in oxygenated artificial cerebrospinal fluid (aCSF; in mM: 126 NaCl, 2.5 KCl, 1 NaH_2_PO_4_, 26.2 NaHCO_3_, 1.3 MgSO_4_, 11 D-glucose, 2.5 CaCl_2_), incubated at 31°C for 30 minutes and then transferring to room temperature for at least another 30 min prior to recordings. Fluorescence was visualized using LED illumination and filter cubes on an Olympus BX51W microscope using a digital camera (DAGE IR-2000). Borosilicate recording electrodes (6-10 MΩ) were used to record neurons. After establishing whole-cell access and after changing the holding potential from -70 to 0 mV cells were given at least 5 min to recover prior to recordings. All recordings were obtained at 31.0°C and acquired using a Multiclamp 700B amplifier and Digidata 1440 digitizer with Axon pClamp 9.0 ClampEx software (Molecular Devices). Data were lowpass filtered at 2 kHz and digitized at 20 kHz. To verify recorded neurons were located within the PNN depleted area brain slices were fixed with 10% formalin for 1h prior to a WFA staining procedure (see above).

For intrinsic excitability studies, a potassium gluconate internal solution was used containing (in mM: 140 K-gluconate, 5 KCl, 10 HEPES, 4 Mg_2_-ATP, 0.5 Na_2_-GTP, 10 Na_2_-phosphocreatine, and 0.2 EGTA). Picrotoxin (100 μM) and NBQX (10 μM) were added to the aCSF to block excitatory and inhibitory synaptic currents. Membrane potential was maintained at -70 mV at rest in current clamp, and cells were presented with a series of current steps (−100 to 200 pA) with a step size of +10 pA. Rheobase was defined as the minimum current step to elicit an action potential. Passive membrane properties were derived using a 10 pA current injection step. For examining the effects of J60 on Gi- and Gq-DREADD transduced neurons, current was injected to elicit basal firing, and J60 (1 μM) was applied after 5 minutes of baseline recording.

For the recording of inhibitory and excitatory postsynaptic currents (IPSC/EPSC), cells were recorded using a Cs-Methanesulfonate (CH_3_CsSO_3_) internal solution containing (in mM: 115 CH_3_CsSO_3_, 10 HEPES, 1 EGTA, 1.5 MgCl, 4 Mg-ATP, 0.3 Na-GTP, 10 Na_2_-phosphocreatine, 2 QX314-Cl, 10 BAPTA-Cs_4_) (Knowland et al., 2017). Spontaneous and evoked EPSCs were obtained at -70mV, and IPSCs were obtained at 0 mV. For evoked recordings a bipolar tungsten stimulation electrode (World Precision Instruments) was used. Series resistance (Rs) was monitored across experiments using a 10 mV pulse, and cells with Rs >20 MΩ or changes in Rs over 15% across an experiment were excluded from analyses. In addition, cells with a holding current below -150 pA were excluded.

Data were analyzed using Clampfit and custom MATLAB scripts. For the quantification of action potential characteristics, the second to fourth action potential waveforms recorded at +80 pA current injection were averaged, filtered using the MATLAB *sgolayfilt* function, and upsampled with the *interp1* fuction using a piecewise cubic hermite interpolating polynomial fit (pchip). Traces were centered at spike threshold (dV/dt > 10 mV/ms) to derive halfwidth, after-hyperpolarization, amplitude and phase data. All other data were analyzed with standard Clampfit functions using unprocessed data except for sEPSC/sIPSC, for which data were preprocessed using a built in 500 Hz Butterworth filter.

### Statistics

All data were analyzed using Prism (v9.0.0; GraphPad). Paired and unpaired student’s t-tests were used for the analysis of cell counts and electrophysiological datasets. Non-normally distributed imaging data and electrophysiological chemogenetic validation studies were examined using Mann-Whitney and Wilcoxon signed-rank tests. Repeated measures analysis of variance (RM-ANOVA) followed by Holm-Šídák post-hoc tests were used for the analysis of behavioral data. Data were screened for outliers using a robust regression and outlier removal (ROUT) algorithm, with a false discovery rate of 1%. Statistical significance was set at 0.05.

## Data and Code Availability

The published article includes all data generated or analyzed during this study. Custom MATLAB code was used to process and analyze electrophysiology data. Code and data are available upon request.

## Acknowledgments

We thank Drs. Jason Aoto, Susan Ingram, and Christopher Ford for original breeder mice used to establish PV-IRES-Cre colonies and for advice on patch-clamp electrophysiology studies. In addition, we thank Katherine Glodoski and Charlie Olson for assistance with immunohistochemistry, and members of the Heinsbroek lab for editing the manuscript and discussion of experiments. This work was supported by National Institutes of Health grants DP5 OD026407 and R01 DA056660.

## Author Contributions

Conceptualization, J.A.H.; Methodology, M.N., G.G., and J.A.H.; Investigation, M.N., G.G., C.N.M., S.D.G., N.K., B.W., N.F., J.A.H.; Resources, J.A.H.; Writing – Original Draft, M.N. and J.A.H.; Writing – Review & Editing, M.N., G.G., C.N.M., S.D.G., N.K., B.W., N.F., J.A.H.;

## Declaration of Interests

The authors declare no competing interests.

---

**Supplementary Figure 1.**
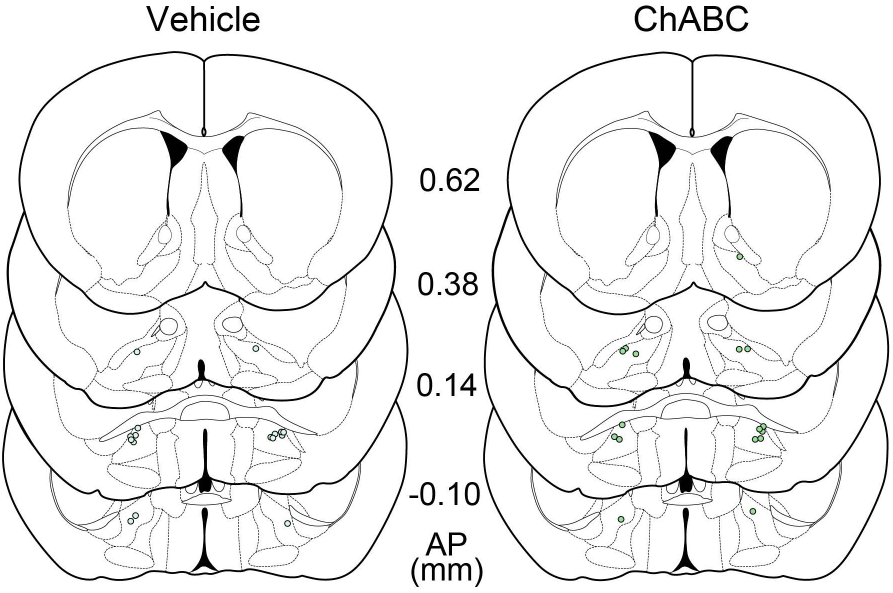
Cannula placements in vehicle and ChABC treated mice. Related to Figure 3. Atlas plates adopted from Paxinos and Franklin (2007) *The mouse brain in stereotaxic coordinates*, 3^rd^ edition show cannula placements in the VP for vehicle (PBS) or ChABC microinfusions in the VP, 24h prior to cue-induced reinstatement testing.

**Supplementary Figure 2.**
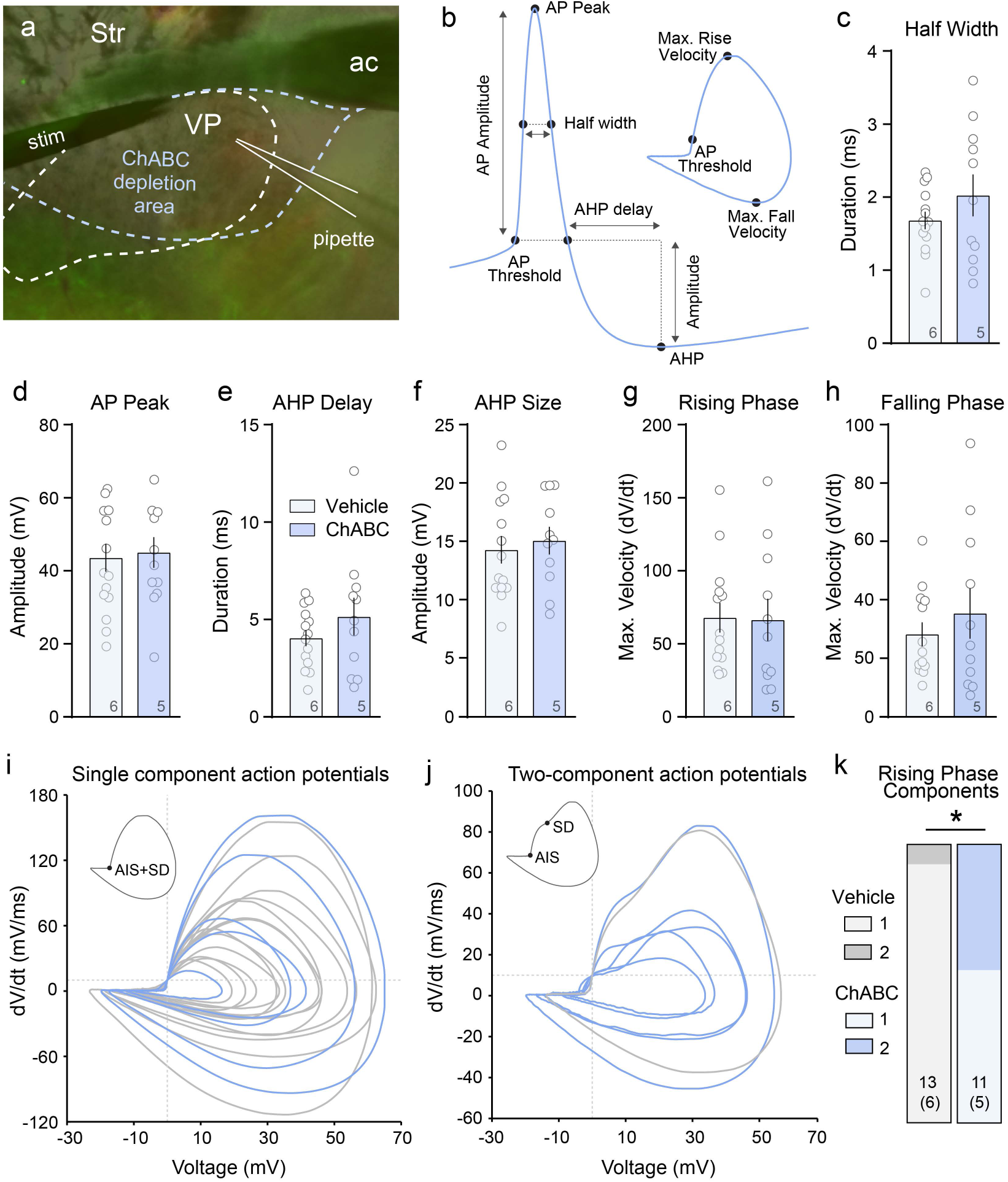
Action potential characteristics following vehicle or ChABC treatment of the VP. Related to Figure 4. (**a**) Representative micrograph showing an overlay of a DIC image of a recorded VP_PV_ neuron with an overlayed image showing tdTomato immuno- and WFA-lectin labeling of the same section to determine whether a cell was recorded within the PNN depletion area (blue dashed line) of the VP (white dashed line) following a ChABC microinfusion. ac = anterior commissure, Str = striatum, stim = bipolar tungsten stimulation electrode, pipette = location of the recording pipette. (**b**) Graphical summary of analyzed action potential characteristics. (**c**) ChABC did not affect the half-width, (**d**) action potential peak, (**e-f**) afterhyperpolarization delay or amplitude, nor (**g-h**) the maximum velocity of the rising or falling phase of the action potential. (**i**) Phase-plane plots showing a single-, or (**j**) two-component rising phase of the action potential broken down by VP_PV_ neurons treated with vehicle (grey) or ChABC (blue). AIS = axon initial segment origin of the action potential, SD = somatodendritic compartment origin of the action potential. (**k**) Quantification of the number of action potentials in each treatment group defined by a single or two-component action potential. * p < 0.05 comparing between vehicle and ChABC. Data shown as mean ± s.e.m.

**Supplementary Figure 3.**
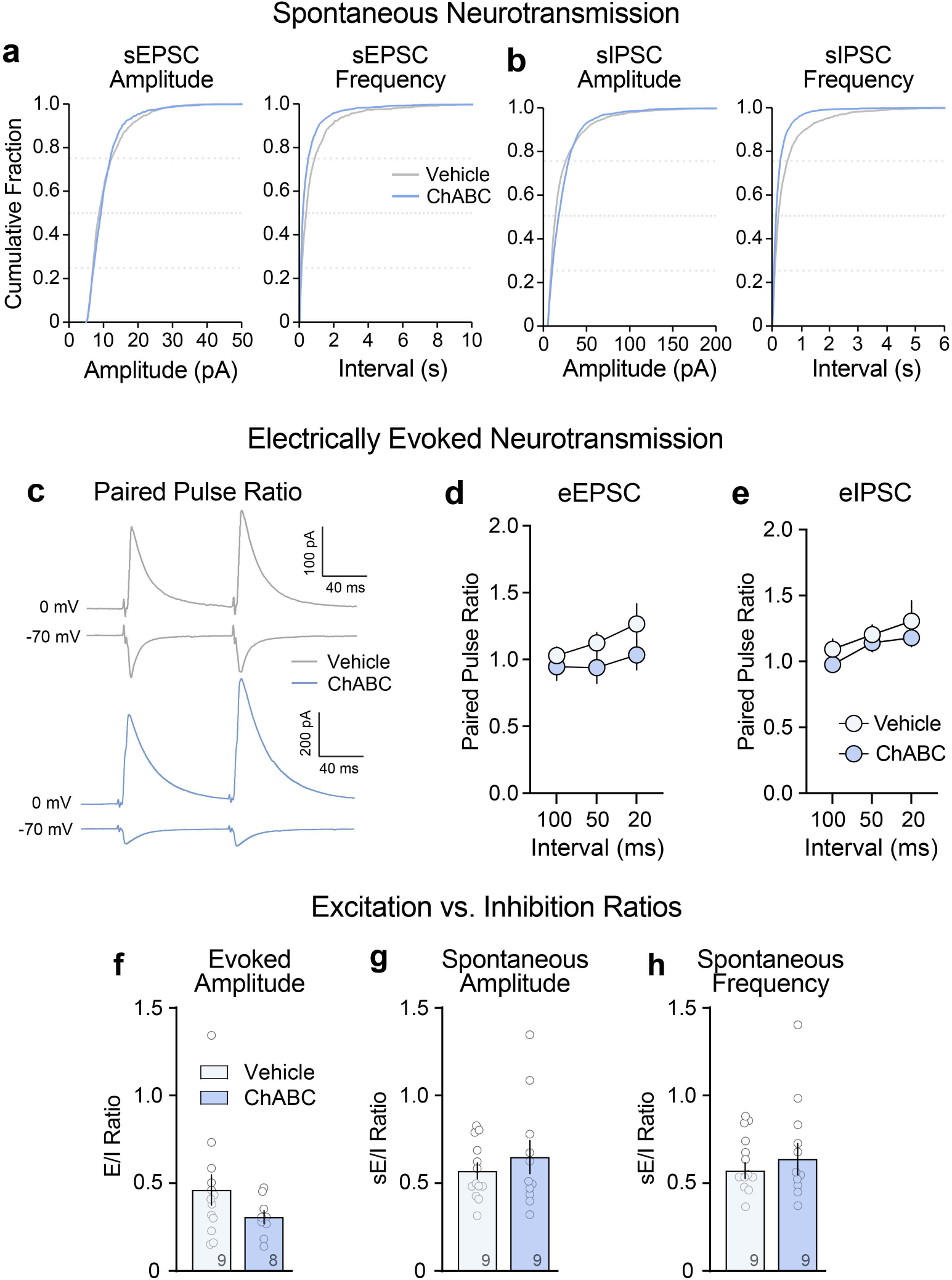
Characterization of synaptic inputs onto VP_PV_ neurons following vehicle or ChABC treatment. Related to Figure 4. (**a**) Cumulative frequency distributions of the amplitudes and inter-event intervals of spontaneous excitatory post-synaptic currents (EPSC) and (**b**) spontaneous inhibitory post-synaptic currents (IPSC). (**c**) Representative traces showing electrically evoked paired-pulse ratios (PPR) after vehicle (grey) or ChABC treatment (blue). EPSCs were recorded at -70 mV and IPSCs were recorded at 0 mV. (**d-e**) Quantification of PPRs after varying inter-pulse intervals show that that ChABC treatment did not affect electrically evoked EPSCs or IPSCs PPRs. (**f**) Electrically evoked EPSC/IPSC ratios did not significantly change following VP PNN depletion. (**g-h**) The ratio of spontaneous EPSC/IPSC amplitude or frequency did not change following ChABC treatment. Symbols in graphs represent individual cells, and numbers in bars represent animal numbers. Data are shown as mean ± s.e.m.

**Supplementary Figure 4.**
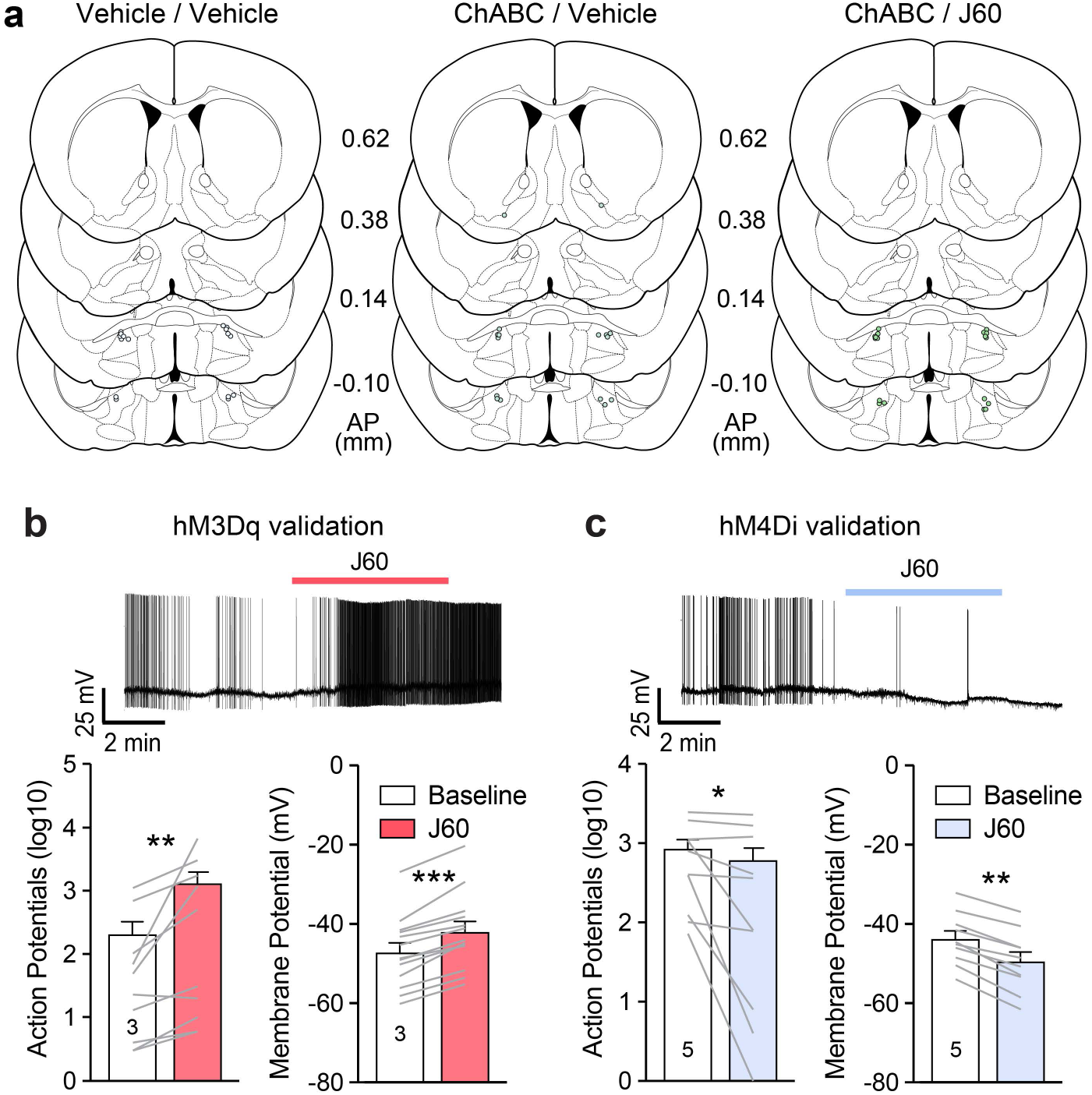
Cannula placements in vehicle and ChABC treated mice and electrophysiological validation of the used chemogenetic strategy for activating or silencing VP_PV_ neurons. Related to Figures 5 and 6. (**a**) Atlas plates adopted from Paxinos and Franklin (2007) *The mouse brain in stereotaxic coordinates*, 3^rd^ edition show cannula placements in the VP for vehicle (PBS) or ChABC microinfusions in the VP, 24h prior to cue-induced reinstatement testing (**b**) Washing J60 onto slices from hM3Dq transduced VP_PV_ neurons results in an increase in spontaneous activity (bottom left), and a depolarization of the membrane potential (bottom right). (c) Washing J60 onto slices from hM4Di transduced VP_PV_ neurons reduces spontaneous activity (bottom left) and hyperpolarizes the membrane potential (bottom right). Lines in bars represent individual cells and numbers in bars represent individual mice. * p < 0.05, ** p < 0.01, *** p < 0.001 comparing between baseline and J60. Data are shown as mean ± s.e.m.

## Notes

### Competing Interest Statement

The authors have declared no competing interest.

